# ISM-FLUX: MINFLUX with an array detector*

**DOI:** 10.1101/2022.04.19.488747

**Authors:** Eli Slenders, Giuseppe Vicidomini

**Affiliations:** Istituto Italiano di Tecnologia, Genova, Italy

**Keywords:** Single-molecule localization, MINFLUX, ISM-FLUX, super-resolution microscopy, single-photon microscopy, single-photon array detector

## Abstract

Single-molecule localization based on the concept of MINFLUX allows for molecular resolution imaging and tracking. However, MINFLUX has a limited field-of-view (FOV) and therefore requires a precise pre-localization step. We propose ISM-FLUX, a localization technique that combines structured illumination with structured detection. We show via simulations that by replacing the point-detector with a small single-photon detector array (e.g., of 5 × 5 elements) and sequentially exciting the sample with four spatially separated doughnut-shaped beams, a localization uncertainty between 1 and 15 nm can be obtained over a FOV of more than 800 nm with 100 photons. The large FOV and the extra spatial information induced by the detector array relax the requirements on prior information on the fluorophore’s position. In addition, ISM-FLUX allows the localization of multiple molecules simultaneously. We calculate the effect of different parameters, such as the relative position of the doughnut beams, the number of detector pixels, the number of photons and the signal-to-background ratio, on the localization uncertainty. We predict that the combination of a good localization precision and the experimental simplicity of ISM-FLUX will help the wide adoption of MINFLUX and other derived microscopy techniques.

## I. INTRODUCTION

The localization of single fluorescent molecules with nanometer precision is important for both single-particle tracking at high spatiotemporal resolution and imaging beyond the diffraction limit. Nowadays, most widely used techniques fall in either of two categories: structured detection (SD) or structured illumination (SI) based localization.

In typical SD based techniques, a camera detector (e.g., a sCMOS or EMCCD) registers the fluorescence collected from a wide-field optical system with a large field-of-view (10-100 *μ*m wide), and the resulting image is analysed to localise the molecule. Most popular implementations of single-molecule localization microscopy (SMLM) techniques for imaging, such as stochastic optical reconstruction microscopy (STORM) [1, 2], photoactivated localization microscopy (PALM) [3, 4], and points accumulation for imaging in nanoscale topography (PAINT) [5] are examples of SD localization. Theoretically, the local<u>iz</u>ation uncertainty of SD techniques scales with 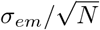, with *σ*_*em*_ the standard deviation of the Gaussian-shaped emission point-spread-function (PSF) on the camera and *N* the number of detected photons. However, other aspects, such as the read-out noise of the camera and different sources of background, negatively affect the precision. As a result, SD techniques typically reach a resolution of about 20-50 nm [6] – although values below 5 nm have been reported [7, 8]. The major advantages of SD techniques are the low-cost implementation – based on a simple wide-field microscope equipped with a sensitive camera – and the possibility to parallelize the localization process. Indeed, a large number of fluorescent molecules can be localized simultaneously, provided that they are farther away from each other than the diffraction limit.

On the other hand, in SI based localization, the information on the position of the fluorophore is encoded entirely in the excitation pattern. By using a laserscanning microscopy architecture, a focused laser beam moves around a single active fluorophore in a (nanometer) precise and pre-defined trajectory, called the targeted coordinate pattern (TCP). A single-element detector, e.g., a single-photon avalanche diode (SPAD) or a photo-multiplier tube (PMT), registers the fluorescence intensity for each position in the TCP and a statistical approach (e.g., maximum likelihood) is used to estimate the position of the molecule. Originally implemented mostly for single-molecule tracking under the general name of real-time single-particle tracking [9, 10], with the introduction of MINFLUX [11], SI based localization became also popular in the context of imaging. MINFLUX showed for the first time the superior localization efficiency when using a doughnut-shaped excitation distribution with minimal intensity in the center. Specifically, in the most typical MINFLUX implementation, the localization uncertainty f<u>or</u> a fluorophore within a circular TCP scales with 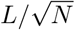, with *L* the diameter of the TCP. Therefore, iteratively reducing the TCP diameter decreases the localization uncertainty more effectively than increasing the number of photons by integrating longer [12]. As a consequence of the *L* tuning, a given uncertainty can be obtained with fewer photons the in conventional SMLM. Similarly to SD techniques, other aspects such as the detector noise and the background have an impact on the effective precision. Importantly, MINFLUX requires a pre-localization of the fluorophore to ensure that the fluorophore is within the initial TCP – a condition that must be met during the whole iterative localization process. Since SI based approaches can make use of a single-photon detector, they can be combined with time-resolved spectroscopy analysis, such as fluorescence lifetime [13]. Other important recent advances in this class of techniques are the possibility to further improve the localization for a certain photon budget by combining it with two-photon excitation (two-photon MINFLUX) [14] or stimulated emission depletion microscopy (MINSTED) [15], and the possibility to be implemented on a commercial laser-scanning microscope by using the conventional bi-dimensional raster scan as TCP (RASTMIN) [16, 17]. However, by spatially integrating all photons with a single-element detector, important information on the molecule’s position is partially lost. This is particularly problematic when the molecule is outside the region spanned by the TCP [12], as the likelihood function may show multiple local maxima far away from the actual fluorophore position. In typical MINFLUX implementations, this lack of robustness demands a precise pre-localization procedure to make sure that the active fluorophore is inside the TCP [18] which, in the case of iterative MINFLUX, may complicate the process.

Here, we propose a molecule localization technique that takes inspiration from MINFLUX and image scanning microscopy (ISM) [19, 20]. Our technique, called ISM-FLUX, combines structured illumination – by means of spatially displaced doughnut-shaped beams that sequentially illuminate the sample with structured detection – by using a camera, i.e., an array of detectors, that records an image of the probed region for each exposure. We calculate the Cramér-Rao bound (CRB) for different experimental parameters, such as the properties of the TCP, the number of photons, the signalto-background ratio, the magnification on the camera, and the number of camera elements to show the effect on the theoretical localization uncertainty that can be expected in each case and to find the optimal experimental settings. We draw particular attention to the case of localizing a fluorophore outside of the TCP, which is the main bottleneck of conventional MINFLUX based techniques that our ISM-FLUX approach can help to overcome. In addition, we show that our technique allows the localization of multiple fluorophores simultaneously. For the sake of completeness, SI and SD localization approaches have been already combined for wide-field SM imaging but with a limited resolution improvement: the typical striped-illumination of a structured-illumination microscope (SIM) can be used to enhance the lateral localization precision by a factor of maximum 2 compared to conventional SMLM [21–23].

## II. MATERIALS AND METHODS

### A Setup

The most versatile implementation of ISM-FLUX can be obtained by replacing the single-element detector of the conventional (non-descanned) MINFLUX setup [11, 12] with a small detector array (e.g., 5 × 5 or 7 × 7 pixels/elements, Fig. 1). Similarly to the single-element detector case, a telescope tunes the effective size of the detector in the specimen plane. An activation laser turns on a single photoswitchable fluorophore in the sample. The activation step can be skipped for imaging methodologies that do not require photo-activation, such as PAINT, or can be replaced with an off-switching procedure, *e*.*g*. for STORM. An excitation laser beam is shaped by a vortex phase mask, or a spatial-light modulator (SLM), forming a doughnut intensity spot in the focal plane of the objective lens. Using an SLM guarantees the additional advantage that the characteristics of the intensity spot can be easily tuned (e.g., switch from Gaussian-to doughnut-shaped). The intensity of the beam is deflected by a pair of electro-optical-deflectors (EODs), such that its central zero is sequentially placed at the four focal plane positions: three equidistant positions on a circle and a fourth position in the center of this circle. A scanning stage to move the sample allows changing the overall position of the TCP. This ability is necessary for sequential localizing different molecules in a large structure, and following a molecule when it moves out of the camera FOV. When a fast update of the TCP position is needed, the stage can be replaced by a galvanometer scanning system [18], although in this case, the array detector must be installed in a de-scanned configuration, decreasing the system’s collection efficiency. Photons emitted by the fluorophore(s) are collected by the objective lens and directed to the detector array by using a dichroic mirror. The four doughnut beams continuously excite the sample until enough photons have been collected for an accurate localization – or until the fluorophore has turned to an off–state. In the case of iterative localization, the TCP can be updated (position and *L* diameter) to improve the localization precision [12]. The detector array is connected to a data acquisition system, both of which need to achieve a high enough temporal resolution to link each photon to one of the excitation patterns.

**FIG. 1.**
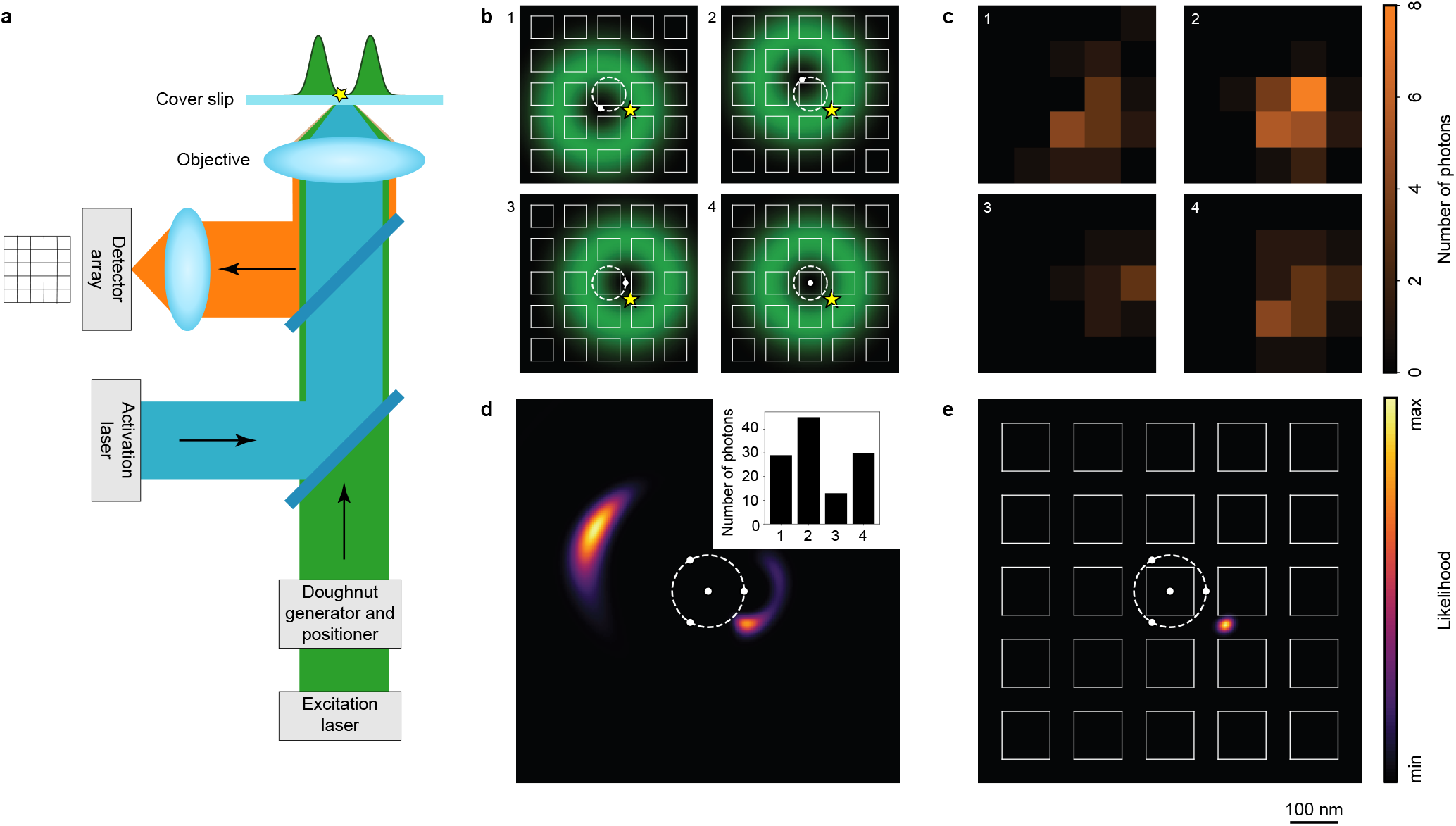
(a) Concept of ISM-FLUX. An activation laser beam activates a single fluorophore in the sample (yellow star) which is sequentially excited (b) by a series of spatially displaced doughnut beams. (c) The fluorescence for each pattern is imaged with a single-photon array detector. (d-e) Corresponding likelihood functions for the localization of the fluorophore taking into account (d) only the fluorescence counts for the different patterns (histogram shown in the inset), *i*.*e*., conventional MINFLUX, and (e) taking advantage of the spatial information of the array detector, *i*.*e*., ISM-FLUX. Simulation settings: 117 emission photons, pixel size and pixel pitch projected in the sample space 100 nm and 150 nm, respectively, *L* = 150 nm, 5 × 5 element detector array. Simulations performed with the help of PyFocus [26].

In stark contrast to the original single-element detector MINFLUX approach, our ISM-FLUX concept does not require the fluorophore to be within the TCP circle for robust localization. The only prior information needed is that one (or more) active fluorophores are present within the detector field-of-view (FOV). This property significantly relaxes the requests for the pre-localization and/or iterative localization process of MINFLUX, thus opening to an alternative implementation with reduced complexity without significantly sacrificing the performance. The edge case is an ISM-FLUX implementation in which the positions of all doughnut beams are predefined and do not have to be updated during the localization experiment, similarly to p-MINFLUX [13]. In this implementation, the TCP sequence can be obtained by splitting the excitation beam into four beams, sequentially turning on each beam with a set of acousto-optic modulators (AOMs) – for example, and recombining them before passing through the vortex-phase plate, or the SLM. A different set of mirrors allows focussing the beams on the sample at the different positions of the TCP. The overall position of the TCP can be changed by means of the scanning stage, or by a galvanometer scanning system when the detector array is used in de-scanned mode. It is worth noting that the setup described above does not provide the possibility to easily tune the *L* parameter. However, since ISM-FLUX can also localize fluorophores outside of the TCP, a small value of *L* can be chosen from the beginning.

A key practical aspect of the ISM-FLUX implementation is the choice of the detector array. A small asynchronous read-out single-photon avalanche diode (SPAD) detector array [24, 25] is a natural choice. They represent the parallelization of the single-element SPAD detector used so far in MINFLUX implementations and the absence of a frame-rate practically does not introduce any new temporal constraints in the implementation. Furthermore, the single-photon timing ability allows a pulsed interleaved excitation (PIE) approach to elegantly implement the TCP sequence [13] (and removing the AOMs), and simultaneously monitoring the fluorescence-lifetime. Correlating the molecule fluorescence lifetime with the position is a key alliance for structural biology applications, and it is not easy to achieve this information with SD based techniques.

However, despite the great momentum that SPAD array detectors experience, state-of-the art detectors show a reduced photon detection probability compared to the single-element SPADs used in the current MINFLUX implementations. Until this gap will be bridged, sCMOS and EM-CCD represent a valuable alternative thanks to their higher quantum efficiency. The prices to pay are the temporal constraints imposed by the frame-rate, the complex combination with single-molecule fluorescence lifetime, and the detrimental effect of the read-out noise. In this context, it is worth mentioning new quantum CMOS cameras with practically negligible read-out noise, thus able to act as a single-photon camera.

### B. Theoretical localization uncertainty

A conventional MINFLUX experiment yields four values, corresponding to the emission photon counts for each of the four positions in the TCP. The fluorophore can be localized with a maximum likelihood estimator, taking into account the combination of the shape and position of the excitation beams with the four photon counts. An ISM-FLUX measurement, on the other hand, does not only provide the number of counts for each position in the TCP but also tells us where in the image plane those photons were detected. ISM-FLUX thus combines the ideas of MINFLUX and conventional single-molecule localization microscopy. As in the application of ISM to improve the resolution of a conventional laser-scanning microscope [20], the detection point-spread-function (PSF) for each element of the detector array is laterally shifted with respect to the detection PSF of the central element. The overall molecule detection function (MDF), which describes the expected number of photons in a pixel as a function of the molecule position, is the product of the excitation intensity (assuming linearity between the laser intensity and the emitted photon count rate (PCR)) and the detection PSF. A total of 100 MDFs can be defined for a combination of four doughnut positions and a 5 × 5 detector array. Note that these 100 MDFs are constant over the experiment and can thus be measured in a reference experiment. Applying the same maximum likelihood procedure as in MINFLUX, but with 100 MDFs instead of 4, yields the fluorophore’s coordinates. In particular, the likelihood function ℒ(**r**_*E*_ **n**) for the fluorophore at position **r**_*E*_ given **n** = (*n*_1_, *n*_2_, *n*_3_, …, *n*_*K*_) photon counts for the *K* = 100 different MDFs can be expressed as

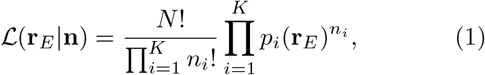

With 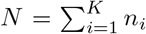 the total number of detected photons and *p*_*i*_(**r**_*E*_) the probability that a photon is detected by MDF_*i*_, *i*.*e*., by one of the 100 combinations of excitation pattern and detector pixel. We assume that both the signal and all background contributions (coming from both out-of-focus fluorescence and detector dark counts) are Poisson distributed and that the background is equal for all pixels and all excitation patterns. Then,

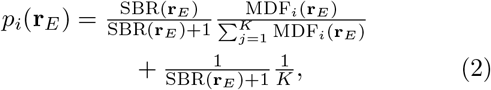

with SBR the signal-to-background ratio. The fluorophore’s position can be estimated as arg max ℒ (**r**_*E*_ ∣**n**) or, equivalently, arg max *𝓁*(**r**_*E*_ ∣ **n**), with *𝓁* = ln() the socalled log–likelihood function.

To obtain an estimate of the precision that can be obtained with this approach, we calculate the Fisher information matrix:

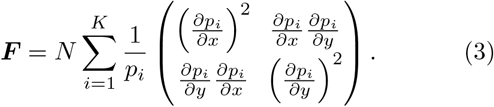

The best obtainable localization uncertainty is given by the Cramér-Rao bound and can be calculated from the inverse of the Fisher matrix, which gives a lower bound for the covariance matrix of the localization uncertainty [11]. Here, we take the arithmetic mean of the eigenvalues of the inverse of the Fisher matrix as a performance metric:

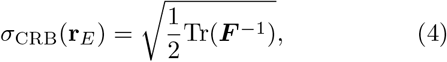

with Tr the trace of the matrix.

### c. Simulation protocol

We assumed an excitation and emission wavelength of respectively 640 nm and 680 nm and an objective NA of 1.4. The pixel size of the simulation space (1024 × 1024 pixels), and hence of all the CRB images, was 2 nm. Unless stated otherwise, each detector element was a square of size 50 μm with a pitch (distance between the centers of two adjacent elements) of 75 μm and the overall magnification was 500×.

The MDFs for the different combinations of excitation pattern and detector element position were simulated in Python. The MDF is *h*_1_ *·* (*h*_2_ *∗ p*), with *h*_1_ the doughnut excitation pattern, obtained with the PyFocus library with the mask set to a vortex phase plate, *h*_2_ the detection (or emission) PSF, calculated with the PyFocus library, *∗* the convolution product and *p* the 2D window function for the corresponding detector element. The different excitation patterns were obtained by shifting *h*_1_ in *x* and *y* to the four positions of the TCP.

We calculated the localization uncertainty *σ*_CRB_(**r**_*E*_)) using Eq. 4 by numerically calculating *p*_*i*_(**r**_*E*_) (Eq. 2) and its derivatives. The total number of photons *N*, and hence the SBR, is position dependent and was calculated as the product of the cumulative excitation intensity at the position of the molecule, limited to a maximum of 1000 counts, and multiplied by the detection efficiency to take into account the effect t hat p hotons e mitted by molecules close to the edge of the FOV have a lower probability of hitting the detector.

For simulating a localization event, for each MDF, photon counts were drawn from a Poissonian distribution with mean *λ* proportional to the respective MDF value at the molecule position. Background counts were taken from a Poissonian distribution with *λ* equal for all MDFs. The vector **n** is the sum of the signal and background counts. To retrieve the position of the molecule, we calculated the likelihood map (Eq. 1) over the whole simulation space and checked for which coordinates ℒ is maximal. To simulate multiplexing, we calculated the signal counts for each molecule as before, summed these counts for each MDF and added the background counts. To retrieve the position of the fluorophores, a least squares optimizer (the ‘SLSQP’ minimizer from the SciPy library in Python) was used to find a (local) minimum in the negative likelihood function, starting at a random point in 4D space. The average distance between the ground truth positions (**q**_1_, **q**_2_) and the retrieved positions (**r**_1_, **r**_2_) was used to quantify the localization uncertainty. Note that the retrieved coordinates for fluorophore 2 may correspond to the ground truth of fluorophore 1 and the other way around, since ℒ (**r**_1_, **r**_2_ ∣ **n**) = (**r**_2_, **r**_1_ ∣**n**). For this reason, we calculated the localization uncertainty for all permutations and took the minimum as the finale result. The same procedure can be used for localizing more than two molecules simultaneously.

Fig. S10 shows a summary sketch of the protocol. All code is available upon request.

## III. RESULTS AND DISCUSSION

### A. Parameters affecting the localization uncertainty

The experimental parameters that the user can control in ISM-FLUX are the optical magnification of the setup (*M*) and hence the region imaged by the detector array, the diameter of the TCP (*L*), the number of signal counts via the excitation power and the pixel dwell time, and the number of pixels of the detector. We calculate *σ*_CRB_(**r**_*E*_) numerically for different values of these parameters and compare the results with conventional “single iteration” MINFLUX, *i*.*e*., MINFLUX with a not iteratively updated TCP.

The extra spatial information in ISM-FLUX has a significant effect on the likelihood function and, consequently, on the CRB, especially when the fluorophore is outside of the TCP circle, Fig. 1 (b-e). In this example, the likelihood function for ISM-FLUX has a single local maximum close to the actual position of the fluorophore. Instead, for MINFLUX, the likelihood function not only has a peak close to the true position, but there is also a large region about 300 nm away from the fluorophore which, in this case, also contains the global maximum. We found similar results for other fluorophore positions outside the TCP circle, Fig. S1.

The difference in the likelihood functions is also reflected in the CRB, Fig. 2. For example, for *L* = 150 nm (*i*.*e*., the same TCP diameter as in Fig. 1), both techniques yield very similar and small CRB values for molecules located close to the TCP center: a precision of about 5 nm can be obtained with 100 photons. In fact, one can show (Supplementary Note 1) that under certain conditions the CRB of ISM-FLUX and the CRB of MINFLUX are identical for a molecule on the optical axis. For molecules located farther from the center, the CRB increases. In the case of MINFLUX, the increase occurs drastically; the average CRB within a field-of-view (FOV) of 300 nm is more than 40 nm, Fig. 2 (b), compared to about 10 nm for ISM-FLUX. The difference increases significantly with increasing FOV, with ISM-FLUX still having a precision below 15 nm for a FOV of 1 μm, while MINFLUX is ineffective for large FOV’s due to regions where the CRB is diverging, Fig. 2 (a). If we consider a more realistic scenario in which the number of signal counts is a function of the fluorophore position, *i*.*e*., the closer the molecule is to the maximum intensity of the doughnut, the more the molecule is excited and the more emission photons will be detected, ISM-FLUX has a very constant CRB up to a FOV of about 600 nm, Fig. 2 (a, last column) and (c). Within this area, the typical increase in localization uncertainty for molecules farther away from the TCP is entirely compensated for by a simultaneous increase in the number of signal counts, Fig. S2. Outside of this area, the signal drops and the CRB increases quickly.

**FIG. 2.**
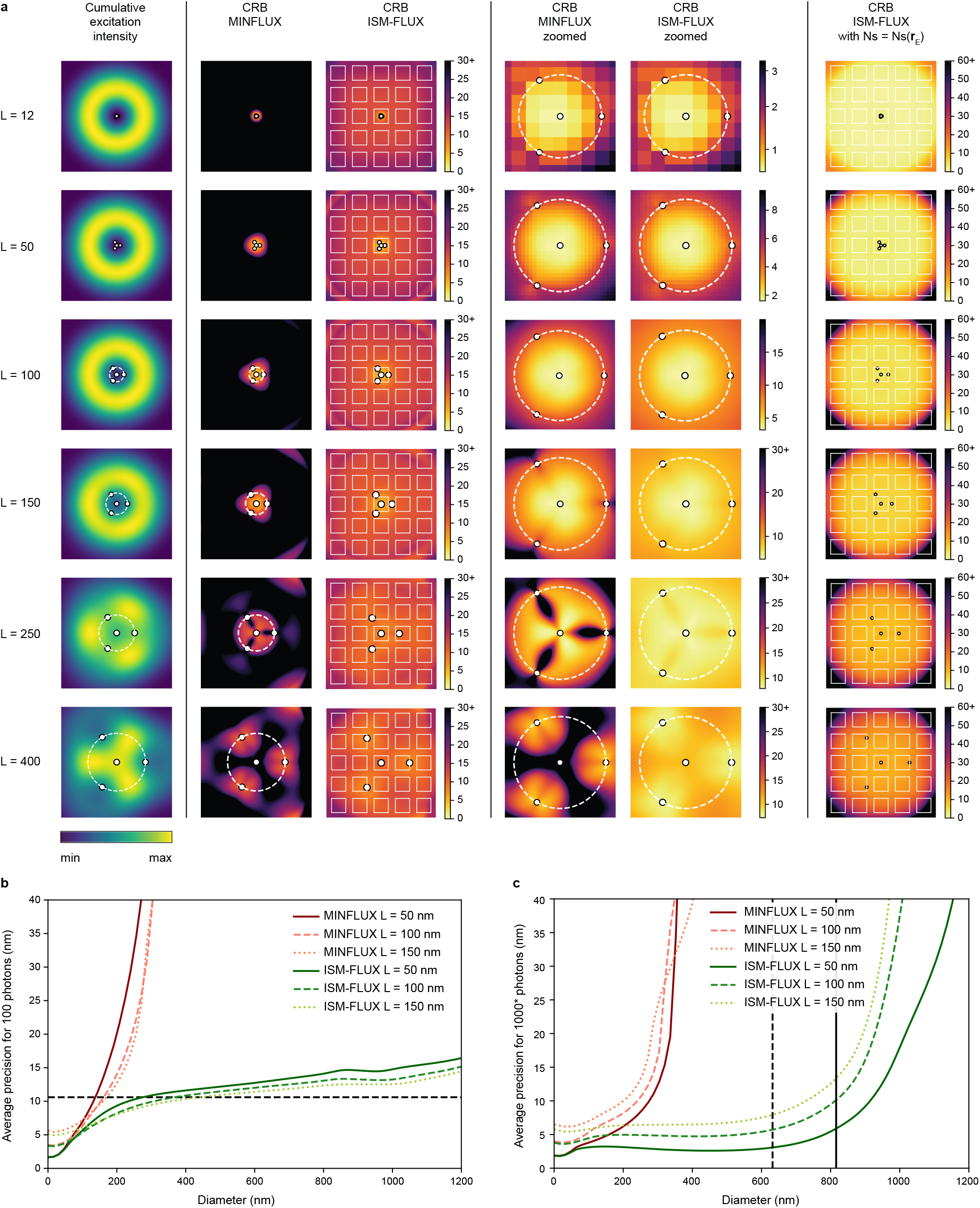
(a) Sum of all four excitation doughnuts (first column) and CRB vs. *L* (other columns), all values in nm. For the last column, the number of signal counts (Ns) was linearly scaled with the local cumulative excitation intensity (with maximum 1000 signal counts in the doughnut maximum) while the number of background counts was assumed to remain constant at 100. A constant 100 signal counts and no background were assumed for the other simulations. The white squares indicate the projection of the array detector in the sample space. 1 pixel is 2 nm. (b-c) Average CRB within a circle centered around the optical axis as a function of the circle diameter for (b) a constant number of photons and (c) a position-dependent Ns, similar to (a, last column), as indicated by the asterisk in the y-axis label. The horizontal line in (b) indicates the theoretical localization uncertainty for standard SMLM in the absence of background, the vertical lines in (c) indicate the region in which approximately 75% and 90% of the excitation occurs.

Clearly, the main advantage of ISM-FLUX is its localization performance for molecules outside of the TCP circle. In fact, the only requirement, and hence the only prior information needed, is that the fluorophore is within the camera FOV and excited strongly enough to produce enough fluorescence photons. This condition is much less strong than in MINFLUX, in which a more accurate prelocalization step is required. Indeed, for a smart choice of the experimental parameters, these requirements are easily fulfilled. For example, when the camera is large with respect to L, the doughnut size, and the emission PSF, any molecule that is excited is also detected by the camera. And given the homogeneity of the CRB over the FOV, no iterative re-centering procedure is needed. Thus, one could simply scan the doughnut beams across the sample until the detected PCR exceeds the background. Then, all of the signal photons can be used for localizing the molecule in a single-step approach. The fact that only little prior information is needed and that the iterative process can be skipped saves time and makes ISM-FLUX more economical on the photon budget and easier to implement.

To find the optimal experimental conditions, we simulated the effect of several parameters on the CRB.

#### 1. Detector size

From the above argument, it is clear that the detector should be large enough to collect as much signal produced by the in-focus plane as possible. The detector size is typically not a tunable parameter, but the magnification with which the emission signal is focused onto the array detector can be adjusted. Within the TCP circle, the magnification has almost no effect on the CRB, Fig. S3 (a-b). A high magnification leads to a homogeneous and good localization precision over the whole detector FOV but at the cost of a smaller FOV, breaking the condition that the detector should cover the excitation region. The CRB is much more heterogeneous for a low magnification, with values ranging from 5 nm close to the optical axis to more than 100 nm near the detector edges. In addition, with a low magnification, the confocal property to block fluorescence from out-of-focus planes is lost, which will lead to an increased background. A good trade-off is M = 500 × (which *e*.*g*. can be achieved with a 100 × objective-tube lens system, followed by an additional 5 × magnification), Fig. S3 (c). This configuration, which is similar to a conventional ISM system, has an average CRB below 7 nm within a FOV of diameter 700 nm for *L* = 100 nm, a maximum of 1000 signal counts in the doughnut maximum, and a constant background of 100 counts.

#### 2. TCP diameter

For small *L* values and for molecules located close to the optical axis, the MINFLUX CRB and the ISM-FLUX CRB are very similar and small, Fig. 2 (a). For *L* = 12 nm, the simulations show a precision of about 0.48 nm with 100 photons for a molecule on the optical axis, in good agreement with the analytically calculated value of 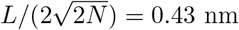 (Supplementary Note 1). The average CRB within the circle of the TCP is less than 3 nm, for both MINFLUX and ISM-FLUX. However, as *L* increases, the homogeneity of the MINFLUX CRB worsens dramatically, while the ISM-FLUX CRB is much less affected and still below 15 nm (within the TCP circle) for *L* = 400 nm. Small *L* values are thus preferable for both techniques, but ISM-FLUX is less prone to the adverse effects that appear at larger *L* values.

#### 3. Number of photons and SBR

SMLM, single-iteration MINFLUX and ISM-FLUX all have a localization uncertainty that scales with 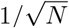, Fig. S4. However, the exact values are different for each technique. The theoretical precis<u>io</u>n in SMLM in the absence of noise is given by 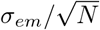, with *σ*_*em*_ the standard deviation of the emission PSF, assumed to be Gaussian. The user has little control over the precision, other than choosing bright fluorophores and sensitive detectors to maximize *N*. In comparison, in SI techniques, also the TCP can be user-defined and influences the precision. Comparing the mean CRB within the TCP circle at high photon counts, ISM-FLUX performs almost two times better than SMLM, and approximately 20% better than MINFLUX. The difference between ISM-FLUX and MINFLUX grows for lower photon counts, arriving at more than 40% below 20 photon counts (assuming 10 background counts), Fig. S4. In addition, unlike MIN-FLUX, ISM-FLUX can also localize molecules with good precision outside of this area, which is a pivotal advantage of ISM-FLUX.

Varying the SBR by varying the number of background counts while keeping the number of signal counts constant, Fig. S5, reveals that the localization precision improves rapidly with increasing SBR. For molecules on the optical axis, a SBR of 1 (both 135 signal and background counts) yields a localization uncertainty that is less than 40% higher than the theoretical limit with zero background. Notably, also the compensation effect in which the higher ‘intrinsic’ localization uncertainty farther away from the optical axis is counterbalanced by a stronger excitation, works better at SBRs around or above 1.

#### 4. Number of pixels

One may expect that increasing the number of detector elements in the array, *i.e.*, for higher K values in Eq. (1-3), either the localization precision or the usable FOV increases, depending on whether, respectively, the detector element size or the overall detector size is kept constant. However, in both cases, the improvements quickly become marginal. Increasing the number of elements while keeping the element size constant, Fig. S6 (a, c), does result in a lower CRB, with the difference increasing with increasing distance from the optical axis. The 3 × 3 array detector has a very limited FOV of about 300 nm. In addition, as pointed out in Section 1, if the detector is much smaller than the excitation and emission area, a significant amount of the fluorescence remains undetected. Assuming a 100 signal counts and an SBR of 20 for the center, for a 5 × 5, 7 × 7, and 9 × 9 array detector, the CRB remains stable around 5 nm up to a FOV of about 600-800 nm, and starts growing rapidly for larger FOVs. This effect, caused by the limited excitation efficiency far away from the optical axis and consequently the low SBR, shows that a 5 × 5 or 7 × 7 array detector is sufficient. Indeed, given that each detector element has a dark count rate, increasing the number of detector elements also decreases the SBR and, hence, can only be beneficial when each element collects a significant number of photons. Increasing the number of detector elements while keeping the overall detector size constant has an even smaller effect on the CRB, Fig. S6 (b, d). By imaging the emission signal with a detector array with more than 5 × 5 pixels, no significant improvement in the obtained localization uncertainty is found. The resulting pixel size is in good agreement with the rule of thumb used in conventional SMLM, i.e., the optimal pixel size is about equal to the standard deviation of the emission PSF [27].

### B. Multiplexing

Apart from the larger FOV, another major advantage of ISM-FLUX is the ability to localize two or more fluorophores simultaneously, which is called multiplexing. The extra spatial information collected by a pixelated detector in combination with a localization uncertainty that is almost homogeneous over the detector FOV allows the localization of multiple fluorophores in a single step. To illustrate multiplexing, we simulated the expected photon counts resulting from two fluorophores at random positions within the detector FOV and performed a maximum-likelihood procedure to retrieve the two fluorophore positions. To localize a single emitter, one can simply calculate ℒ(**r**_*E*_ ∣ **n**) for all reasonable emitter positions and check where ℒ reaches its maximum. This brute-force approach is not sustainable for localizing multiple fluorophores. For example, for two fluorophores, the likelihood function spans a 4D space ℒ (**r**_*E*,1_, **r**_*E*,2_ ∣ **n**), with **r**_*E*,1_ and **r**_*E*,2_ the (x,y) positions of the two fluorophores. Calculating ℒ for all possible combinations of the two positions would require an infeasible amount of computer memory and CPU time. Instead, we used an iterative optimization algorithm to find a local maximum of, ℒ starting at random positions of the fluorophores.

Fig. 3 shows an example of the simultaneous localization of two molecules. Although both molecules emit a significantly different number of photons (220 *vs*. 131) and the overall SBR is relatively low (3.5), both molecules can be localized with similar precision. Repeating the simulation 50 times, we found a median set of coordinates for both fluorophores that was equal to the ground truth, with a localization uncertainty (calculated as the median distance between the retrieved coordinates and the ground truth) of 14 nm for the particle closest to the doughnut center and 16 nm for the other particle. In general, Fig. 3 (d), we found that for two fluorophores at random positions within the excitation region, in 50% of the cases both molecules can be localized with a precision of less than 12 nm, assuming that the molecules are at least half a Rayleigh distance apart. Given the diffraction limit, one cannot expect to distinguish multiple simultaneously active molecules that are within a much shorter distance.

**FIG. 3.**
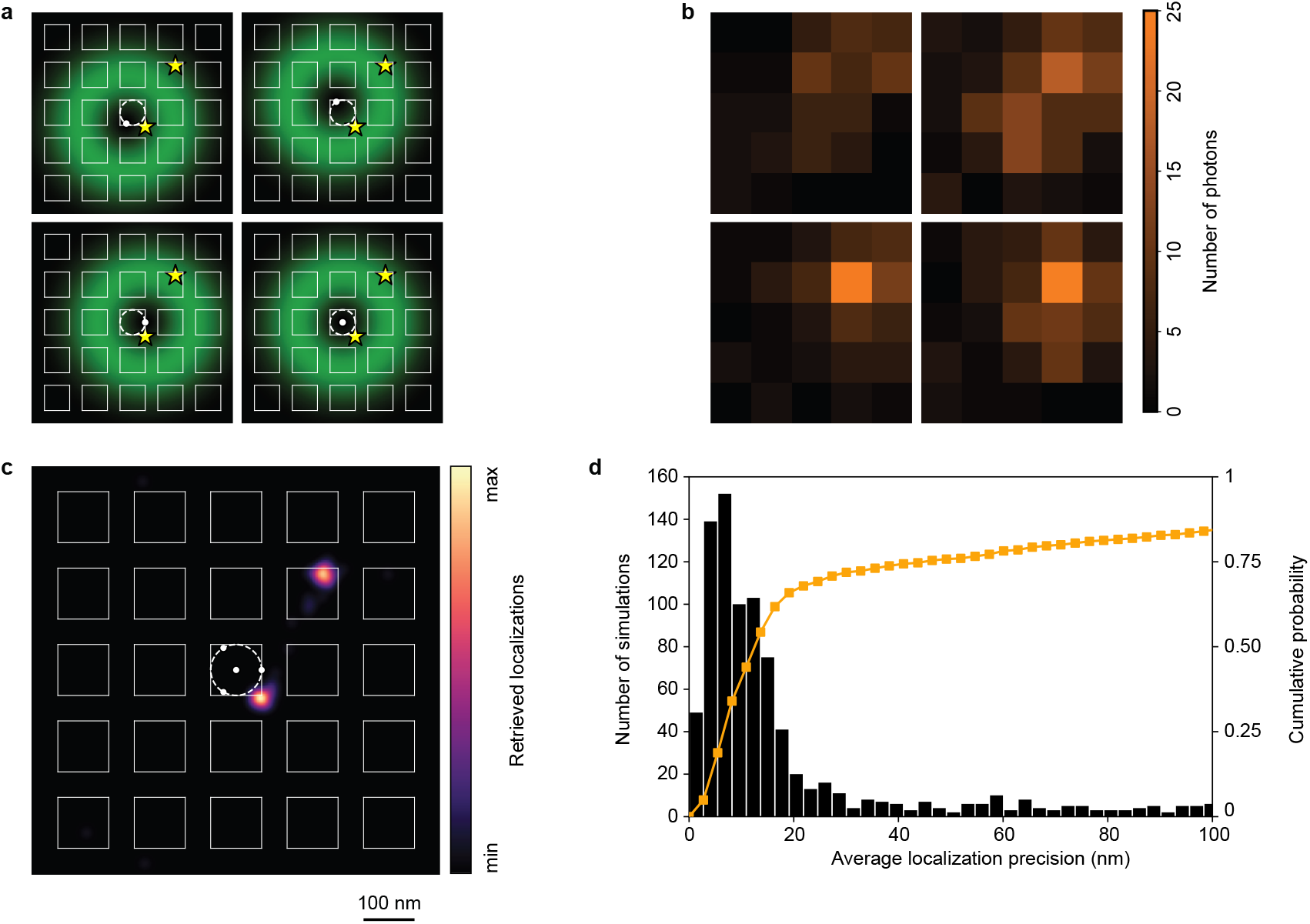
Multiplexing with ISM-FLUX. (a) Example of multiplexing with two molecules located at the positions indicated by the stars. (b) Corresponding observed photon counts for a single simulation. Simulation settings: L = 100 nm, in total 351 signal counts, SBR 3.5. (c) Resulting image for 50 simulations: for each molecule, the retrieved locations were convolved with a Gaussian with standard deviation 9.3 nm – corresponding to the estimated CRB for a single particle under the same conditions – and summed over all simulations. Fig. S7 shows the raw localizations retrieved. (d) Histogram of the average precision for 1000 simulations of two molecules located at random positions within the FOV, at a distance larger than half the Rayleigh limit. Same simulation settings as in (b), with the number of emitted photons linearly scaled with the cumulative excitation intensity. In color the cumulative probability. In 50% of the cases, a localization uncertainty of less than 12 nm is found. The localization uncertainty is defined as the average of the two distances between the retrieved molecule positions and the ground truth positions.

### G. General discussion and detector choice

Localizing a fluorophore with MINFLUX based techniques typically consists of two steps. In the first step, either the fluorescence intensity is measured at the different TCP positions with a regularly Gaussian-shaped intensity distribution or a camera image is made under wide-field illumination. The resulting data gives a first approximation of the fluorophore position. In the second step, the MINFLUX process with a doughnut intensity profile is performed starting at the previously estimated position and - in the case of iterative MINFLUX - decreasing *L* in each iteration. This approach has multiple limitations. Firstly, pre-localization procedures that make use of a regular beam require a setup in which the intensity focal distribution can switch from a Gausian-to doughnut-shaped beand back, which adds experimental complexity to the technique. Secondly, the iterative approach also adds computational and experimental complexity, as the molecule position has to be estimated in real-time, and the TCP radius needs to be updated continuously and in real-time. Here, we presented ISM-FLUX, which requires a less strict pre-localization step – that can be done with the same array detector as the actual localization, and which eliminates the need for a fast iteratively updated TCP. In fact, the iterative approach can be completely omitted in ISM-FLUX. By replacing the point-detector with a small pixelated detector, a molecule can be localized over a large FOV, removing the condition that the molecule has to be within the TCP circle. For typical experimental parameters, ISM-FLUX has a homogeneous localization uncertainty over a 600 nm FOV. In other words, if the molecule is too far from the optical axis to be localized, it is also too far from the doughnut beams to be excited in the first place. This large FOV effectively relaxes the temporal condition for updating the FOV position in the case of the application of our ISM-FLUX principle to real-time single-molecule tracking. Similarly to MINFLUX, the CRB decreases with decreasing *L* and increasing *N*. Our simulations show that the detector array should ideally be 5 × 5 or 7 × 7 pixels. Fewer pixels will lead to a less precise localization; more pixels will not significantly enhance the theoretical performance and will increase the dark count rate.

Although many structured detectors can provide the spatial information needed to perform ISM-FLUX, asynchronous read-out SPAD array detectors [24, 28] are a very good candidate for the experimental realization. These detectors have single-photon sensitivity, no read-out noise, and a high photon timing precision (in the range of hundreds of picoseconds), which is necessary to link each photon to one of the excitation doughnuts when PIE is used. The fact that only a small number of detector elements is needed, thus a relatively low amount of data needs to be transferred/stored by the acquisition system, allows photon time-tagging recording mode [29]. Having access to the nanosecond-resolution arrival times makes the combination of ISM-FLUX with lifetime feasible, which can give extra information on the macro-molecular structure or its microenvironment and allows for lifetime-based multi-species detection [30, 31]. In the context of single-particle tracking, the high timing precision of SPADs may be exploited to study microsecond-scale dynamics with ISM-FLUX.

Fig. 4 shows a comparison between MINFLUX and ISM-FLUX for a realistic set of detector parameters. The development of single-photon (array) detectors is a continuously evolving field, with new detectors with better specifications (in terms of quantum detection efficiency, fill factor, dead time, cross-talk, dark count rate etc.) appearing regularly. A state-of-the-art single-element detector used in confocal microscopy has a quantum yield of around 65% at 650 nm. In comparison, the most recent SPAD array detectors [24, 32] have a quantum detection efficiency around 40% at this wavelength. Assuming that a microlens array is installed in front of the detector to maximize the fill factor to 78.5%, the overall photon detection probability (PDP) is 31.4%, almost two times lower than for a single-element detector. In addition, the dark count rate (DCR) – and hence the background – is typically higher. While a single-element detector may have a DCR around 250 Hz, for a 25-element SPAD array detector, this may be up to 2500 Hz for all elements combined, i.e., one order of magnitude worse. Here, we simulated the CRB for MINFLUX with 128 photons and an SBR of 5 (in the center, photon counts scaled with the excitation intensity for off-center positions) and ISM-FLUX with 62 photons and an SBR of 0.5. Molecules close to the optical axis can be localized with about 2.8 nm and 7.4 nm precision for MINFLUX and ISM-FLUX, respectively. Clearly, the better detection efficiency and SBR in MINFLUX are advantageous. Fig. S8 compares in more detail the CRB for array detectors and single-element detectors. For all values of *L*, an array detector outperforms a single-element detector with the same specifications in terms of PDP and DCR, both for locating molecules close to the optical axis and outside of the TCP. However, current state-of-the art SPAD array detectors do not reach this ideal situation due to a limited fill factor and higher DCRs. As a result, their performance in locating molecules close to the optical axis may actually be worse than with a single-element detector, i.e., the coordinates in Fig. S8 may lie below the line of equal performance. However, a key advantage of ISM-FLUX is the very stable localization uncertainty up to more than 600 nm FOV while for MINFLUX the CRB increases with more than one order of magnitude over the same FOV, as shown in the right panel of Fig. 4 and the right panels of Fig. S8.

**FIG. 4.**
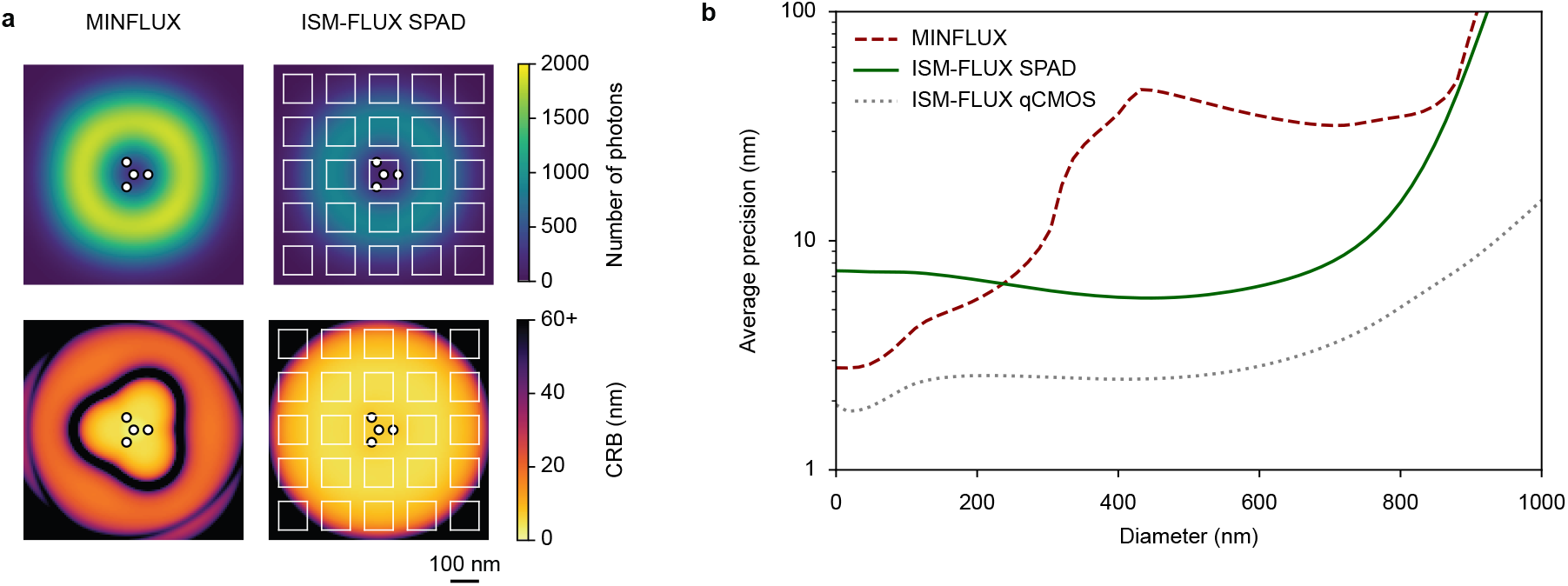
Comparison between MINFLUX and ISM-FLUX for realistic detector parameters. (a) Total number of detected photons as a function of the molecule position (top) and the CRB (bottom). On the optical axis, the SBR is 5 for MINFLUX and 0.5 for ISM-FLUX. L = 100 nm. (b) Average localization uncertainty within a circle around the optical axis as a function of the diameter of the circle. *ISM-FLUX qCMOS* refers to a detector with 90% PDP and zero noise.

Instead of SPAD array detectors, one could also implement ISM-FLUX with other types of detectors, such as sCMOS camera’s. For example, the ORCA-Quest qCMOS camera (Hamamatsu Photonics, Japan) has a peak quantum efficiency of 90% and close-to-zero read-out noise. Such a detector would not only perform better than current SPAD array detectors but also outperform typical single-element detectors. Fig. 4 shows the precision that can be obtained with such a detector in the absence of any noise, which, clearly, is an optimistic scenario. In addition, the qCMOS detector has a standard frame rate of 120 Hz, which is too slow to allow the combination of ISM-FLUX with the measurement of the fluorescence lifetime.

## IV. CONCLUSIONS

We predict that the main advantage of ISM-FLUX will be the simplicity of the experimental implementation. Indeed, our simulations show that current SPAD based ISM setups [20, 28, 33] require only minor modifications for ISM-FLUX measurements. On the excitation side, the conventional laser line can be replaced with four lines in PIE mode, focused at the four positions of the TCP. On the detection side, it is sufficient to update the data acquisition protocol such that each photon can be linked to one of the doughnut beams [13]. The magnification on the detector can be the same as in ISM, which corresponds to about 1 to 1.5 Airy units for each side of the detector. In addition to a good localization precision and large FOV, this magnification also has the optical sectioning effect, which is important for 3D samples or for keeping the background low in samples with DNA-PAINT based blinking [34]. Notably, the setup simplifies further when extending our ISM-based approach to the RAST-MIN architecture. Here, the doughnut-shaped excitation pattern is raster-scanned by the galvo-mirror, making the four beam splitting not necessary. We thereby envisage another SI based localization approach with a further simplified setup. Finally, in the context of imaging, by removing the condition that the molecule must be within the TCP circle, we showed that ISM-FLUX allows fast imaging, since multiple active molecules in the FOV can be localized simultaneously. Here, (i) we assumed that the number of active molecules was known *a priori* and (ii) we used an updated likelihood protocol in which the maximum was found with an iterative optimization algorithm. Regarding the first aspect, whether one or more molecules are active at any given time is unknown, but this information can be derived from the observed photon count rate or through photon antibunching analysis. We demonstrate the first approach in Fig. S9. Alternatively, in antibunching analysis, the almost null tendency of photons produced by a single emitter to arrive in pairs is exploited, leading to a dip in the second-order correlation function of the photon arrival times in the case of a single emitter. SPAD array detectors provide the necessary features for the implementation of this quantum imaging technique [35]. Regarding the second aspect, the analysis protocol used here is based on a standard iterative least squares optimization function. Although we show that the median retrieved positions coincide with the ground truth, there are outliers. It is worth checking whether different optimization algorithms perform better. This will be especially important in more complex cases, such as the localization of more than two emitters simultaneously.

Although here we show only 2D localization, ISM-FLUX may take inspiration from both MINFLUX and conventional SMLM to be extended to 3D. In MINFLUX, a combination of a vortex and a top-hat phase mask in the excitation beam path can be used to create a 3D doughnut [12]. The beam can be focused at different z positions, for example with an electrically tunable lens, to extend the TCP to 3D. Alternatively, the z-position can be controlled with a deformable mirror that adds defocus to the excitation beams [18]. In the case of 3D PIE, both approaches would be too slow, but since the four laser beams are separately accessible, they can be made slightly diverging or converging to control the z position. Similar to conventional SMLM, also adding astigmatism by placing a cylindrical lens in the emission beam path allows for 3D localization. In all cases, the same localization procedure can be used as for 2D, but with the MDFs now extended to 3D functions.

In summary, ISM-FLUX provides an elegant solution to some of the experimental challenges of MIN-FLUX based techniques while maintaining the nanometer scale spatial resolution and the combination with time-resolved imaging. We are convinced that these advantages will trigger many laboratories to develop ISM-FLUX microscopes and help localization microscopy with nanometer precision become routine.

## ACKNOWLEDGMENTS

The authors thank Dr. Giorgio Tortarolo, Dr. Alessandro Zunino, Andrea Bucci, Francesco Fersini, and Simone Civita for the useful discussions, and Fernando Caprile and Dr. Luciano Masullo for their kind assistance with the use of the Python package PyFocus. This research was supported by the European Research Council, Bright Eyes no. 818699 (G.V.) and by the European Union’s Horizon 2020 research and innovation programme under the Marie Sk1odowska-Curie grant agreement no. 890923 (SMSPAD) (E.S.).

## SUPPORTING INFORMATION

### Supplementary Note 1: analytical calculation Cramér-Rao bound

We calculate the CR bound analytically for a molecule close to the center of the detector, i.e. (*x*_0_, *y*_0_) → (0, 0). We define the doughnut beams at positions (*p*_*i*_, *q*_*i*_) (i = 1,…,K) as the product of a second order polynomial with a Gaussian. Then, for doughnuts close to the optical axis ((*p*_*i*_, *q*_*i*_) → (0, 0)), the excitation intensity is:

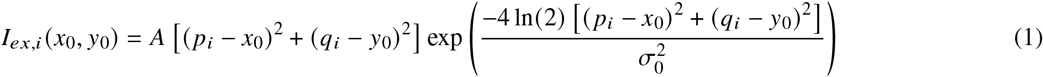

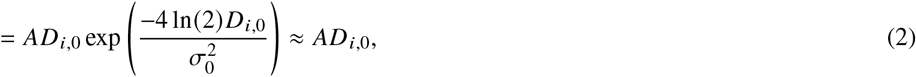

with *A* a scaling factor, 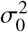 describing the doughnut diameter, and *D*_*i*,0_ the shorthand notation for the squared distance between the doughnut *i* and the molecule position: (*p*_*i*_ − *x*_0_)^2^ + (*q*_*i*_ − *y*_0_)^2^.

We approximate the emission PSF by a 2*D* Gaussian with standard deviation σ.

Under the assumption that the detected fluorescence intensity is proportional to the excitation intensity, the expected relative number of photons that will be detected by each pixel of the array detector under illumination of doughnut *i* (*p*_*i*_) is proportional to the product of the excitation intensity and the emission PSF centered around position (*x*_0_, *y*_0_):

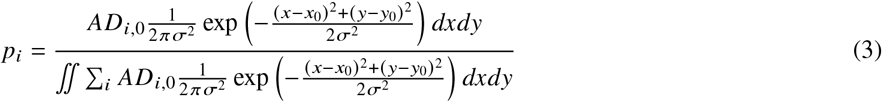

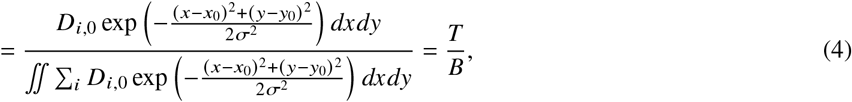

with *T* the numerator and *B* the denominator. We assume an infinitely large detector (which is reasonable if the detector is larger than the emission PSF and the molecule is close to the optical axis) with infinitely small pixels. We omit the factor *dxdy* in *T* to work with photon *densities*.

To calculate the CRB, we need the partial derivatives of *p*_*i*_ to *x*_0_ and *y*_0_. We first calculate the derivatives of *T* and *B*.

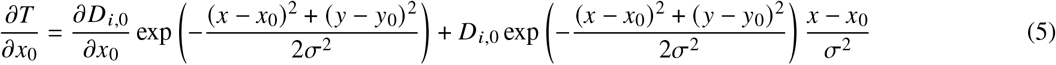

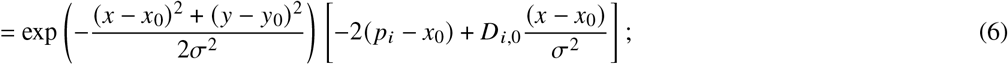

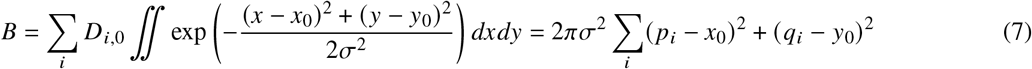

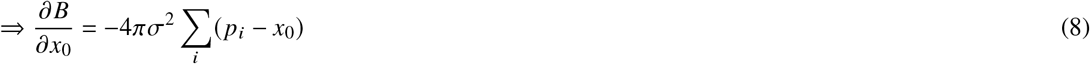

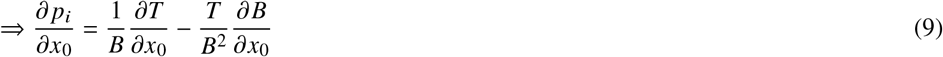

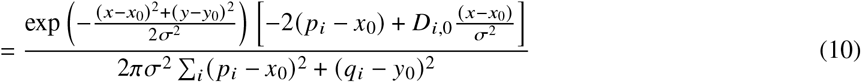

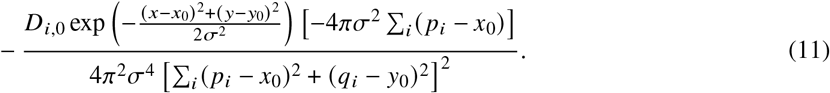

For a molecule on the optical axis, we have (*x*_0_, *y*_0_) = 0, 0 and Σ_*i*_ (*p*_*i*_ *x*_0_) = Σ_*i*_ *p*_*i*_ = 0 for a symmetric TCP. Computing 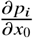, we find (and for symmetry reasons also for 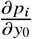):

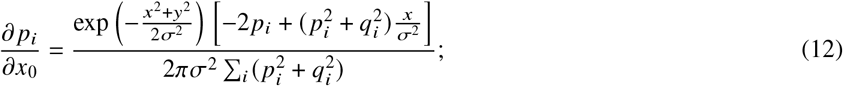

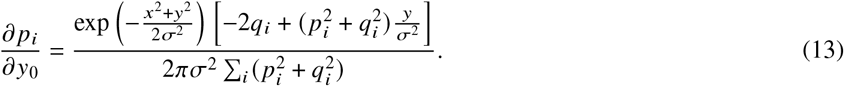

Next, we calculate the following three terms for a molecule on the optical axis 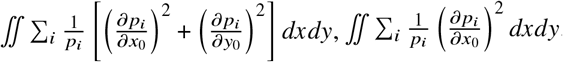, and 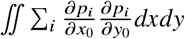

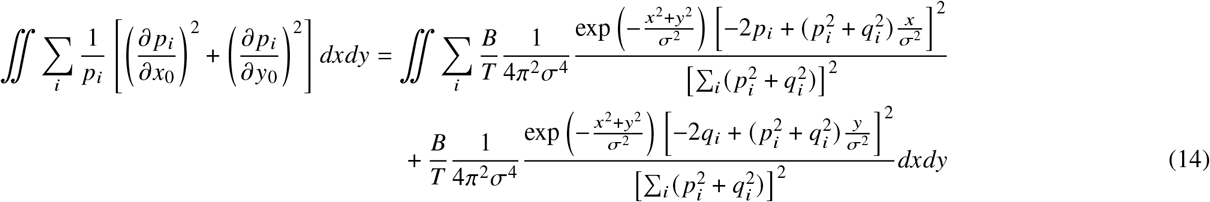

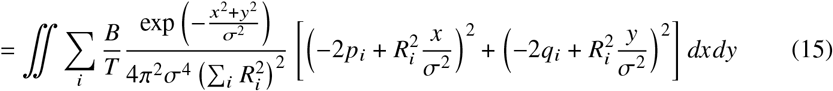

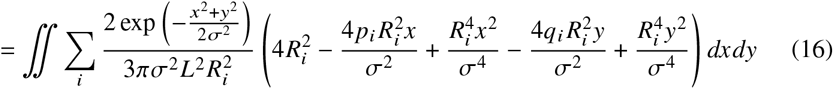

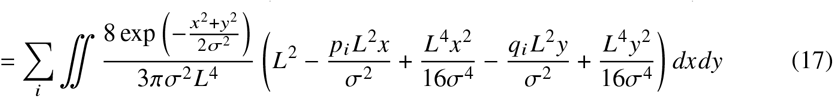

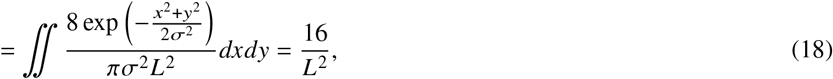

where 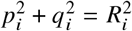. We made use of the fact that the integral of an odd function over R is 0 (if finite), the TCP pattern consists of t hree pos itions uniformly distributed on a circle with radius *L*/2 centered around the optical axis, 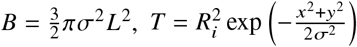, and *L* ≪ σ. We apply the same assumptions to the other terms:

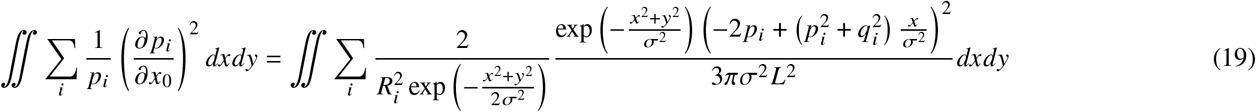

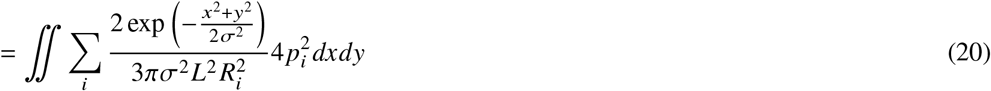

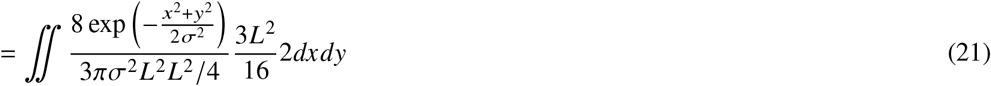

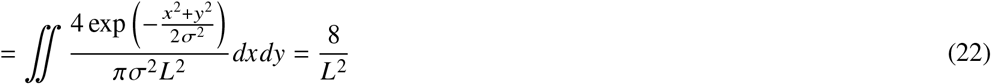

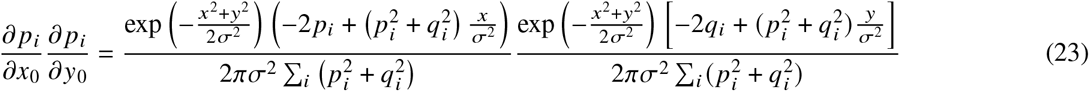

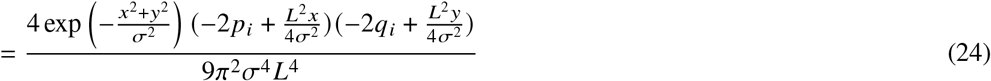

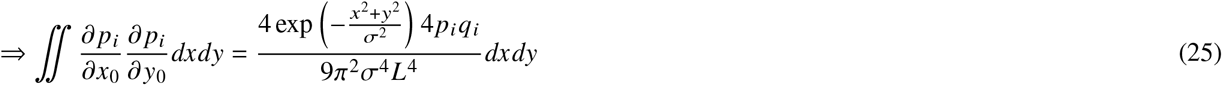

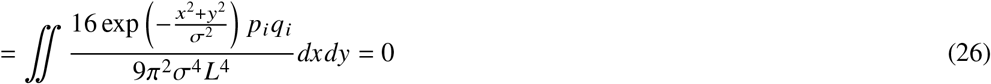

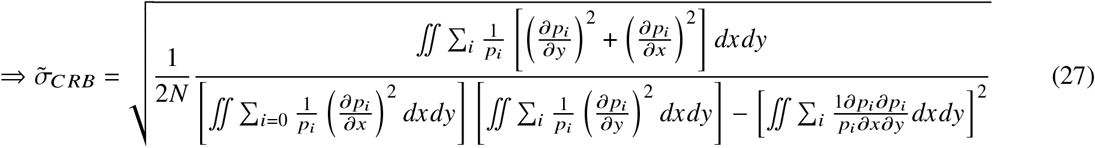

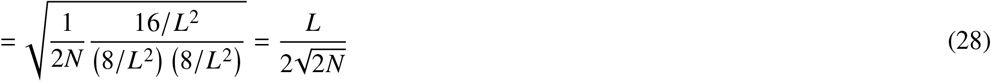

**Figure S1:**
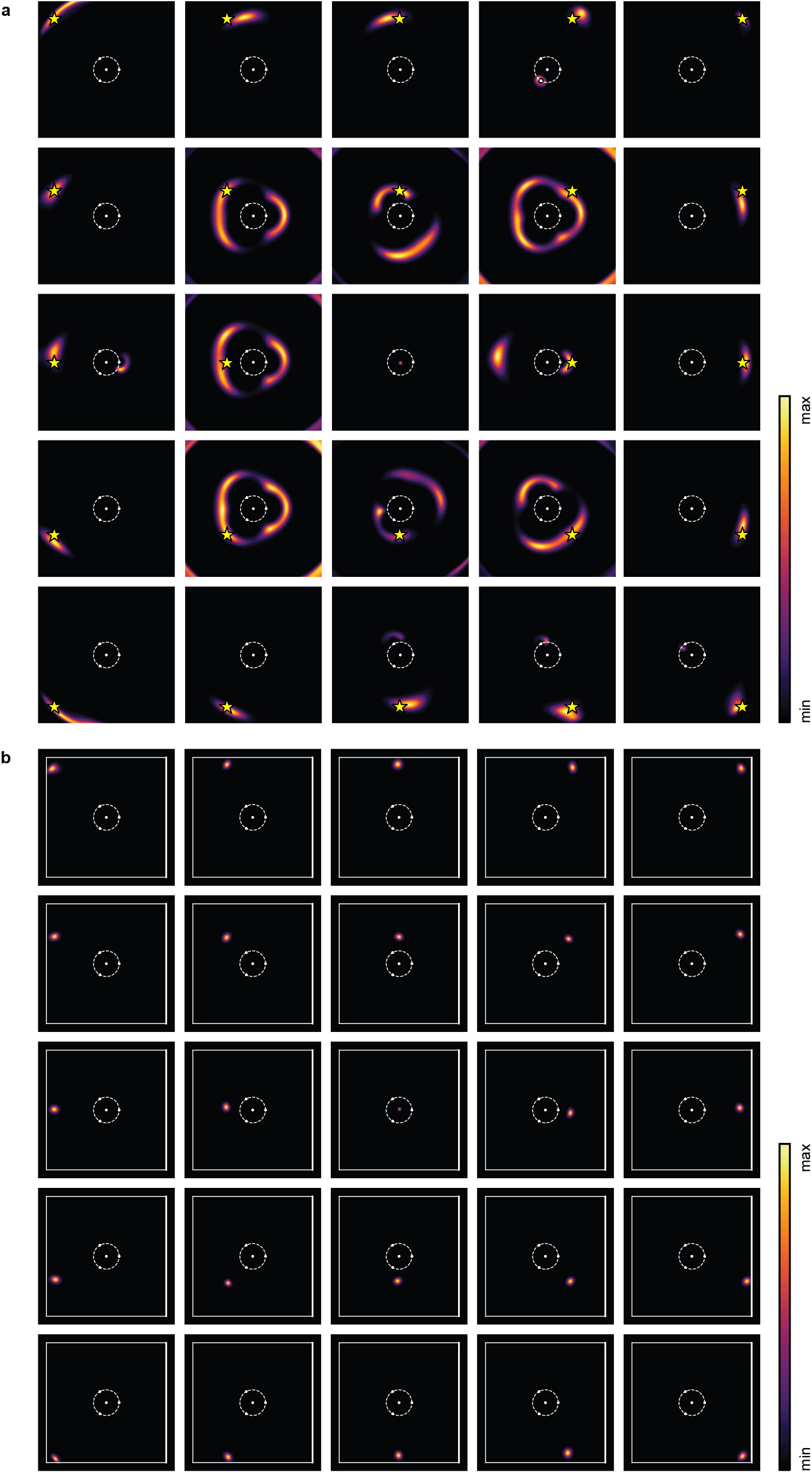
Simulated likelihood functions for (a) MINFLUX and (b) ISM-FLUX for different positions of the fluorophore (yellow star). Simulation settings: 100 signal counts, no background. The circle indicates the TCP, *L* = 150 nm. For visualization purposes, the fluorophores are not shown in all plots. All images show a FOV of 800 nm.

**Figure S2:**
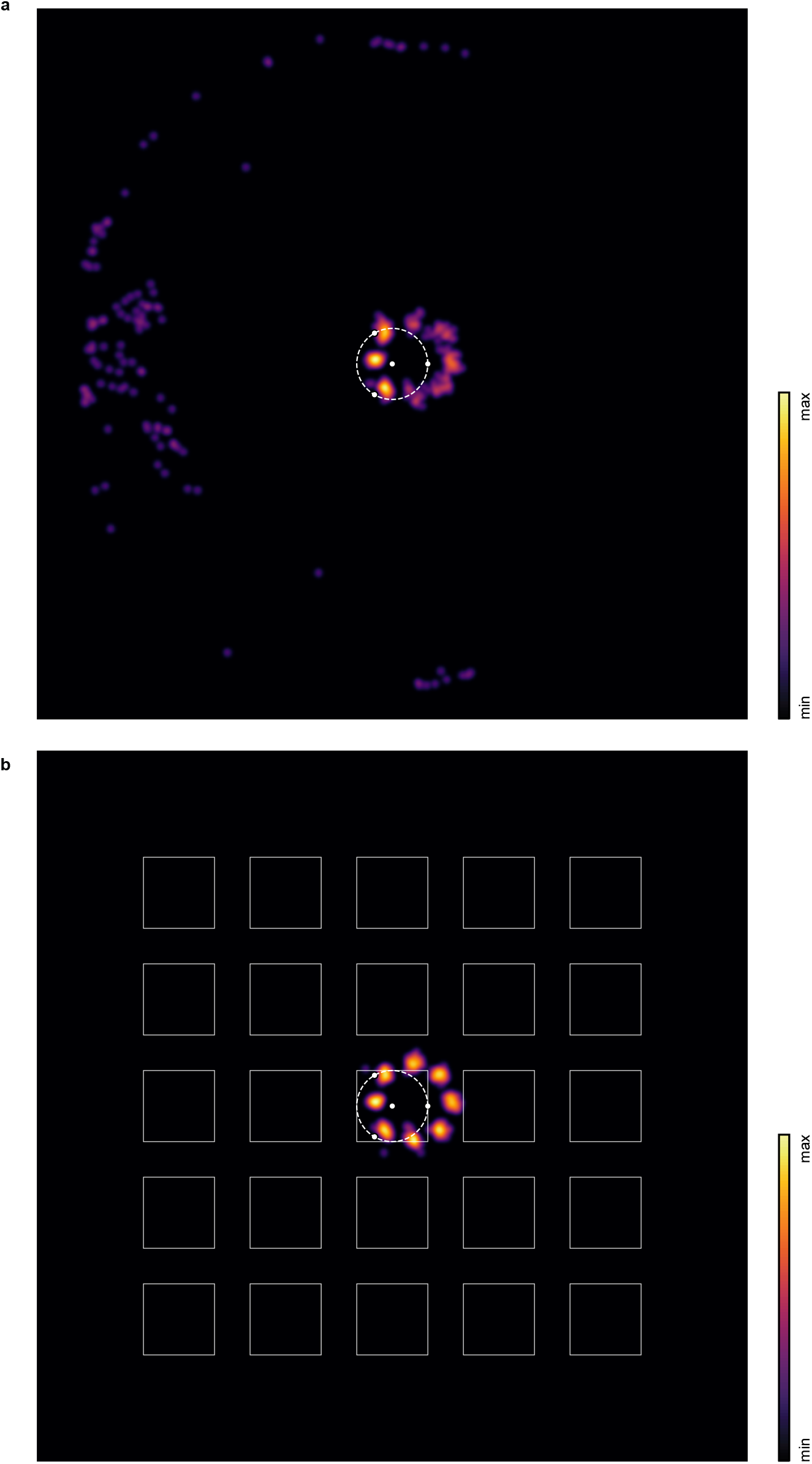
Simulated images of a nuclear pore complex, consisting of eight units located in a regular pattern on a circle of diameter 110 nm with (a) a single-element detector and (b) an array detector. The TCP is kept fixed for all localizations, centered around the optical axis with *L* = 100 nm. Sum over 50 localizations per molecule, each localization performed with between 125 photons (close to the optical axis) and 376 photons (farther away from the optical axis), photon counts linearly scaled with the excitation intensity, no background assumed. Localizations convolved with a 2D Gaussian with *σ* = 2 pixels (4 nm). Logarithmic color map.

**Figure S3:**
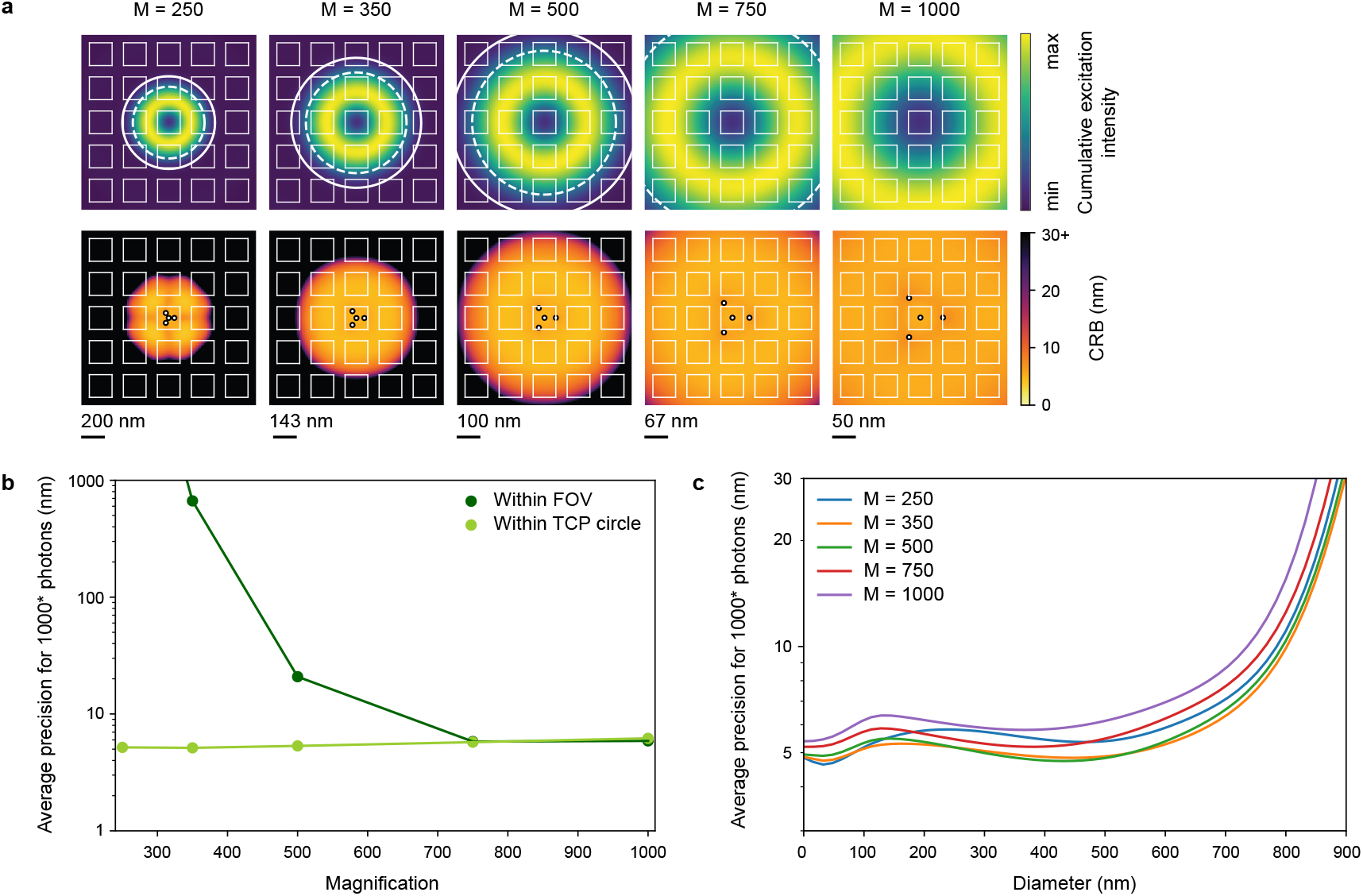
(a) Cumulative excitation intensity (top) and ISM-FLUX CRB (bottom) as a function of the system magnification. The dashed and continuous circles indicate the regions in which respectively 75% and 90% of the excitation intensity occurs. (b) Average CRB within the TCP circle and detector FOV as a function of the magnification. (c) Average CRB within a circle of a given diameter as a function of the diameter. Simulation settings: *L* = 100 nm, SBR linearly scaled with the overall excitation intensity (denoted with the asterisk in the y-axis label), with maximum 1000 emitted photons and a SBR of 10 in the doughnut maximum.

**Figure S4:**
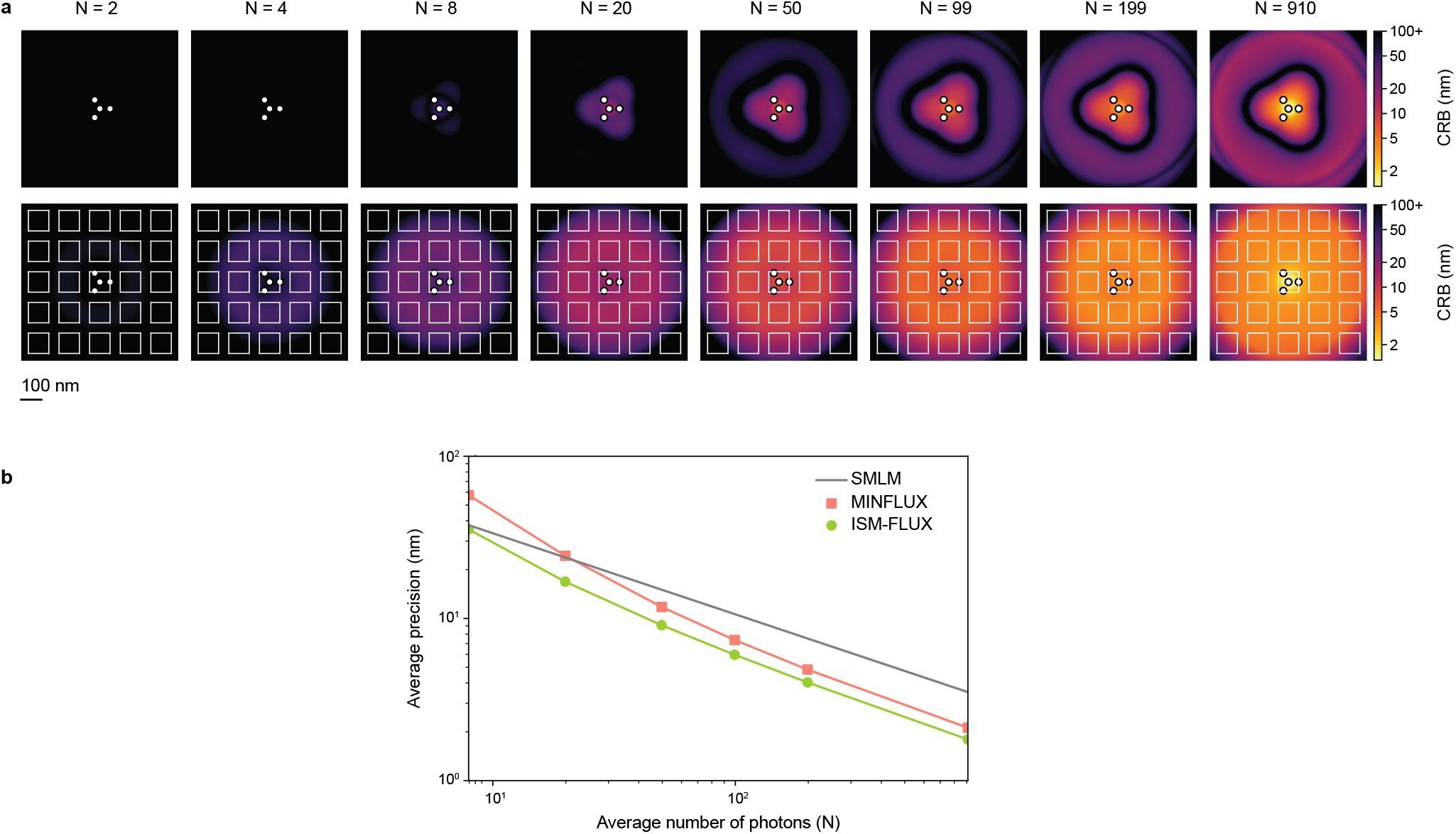
(a) MINFLUX CRB (top) and ISM-FLUX CRB (bottom) as a function of the average number of detected photons within the TCP (N). (b) Average CRB within the TCP circle. Simulation settings: *L* = 100 nm, SBR linearly scaled with the overall excitation intensity, 10 background counts.

**Figure S5:**
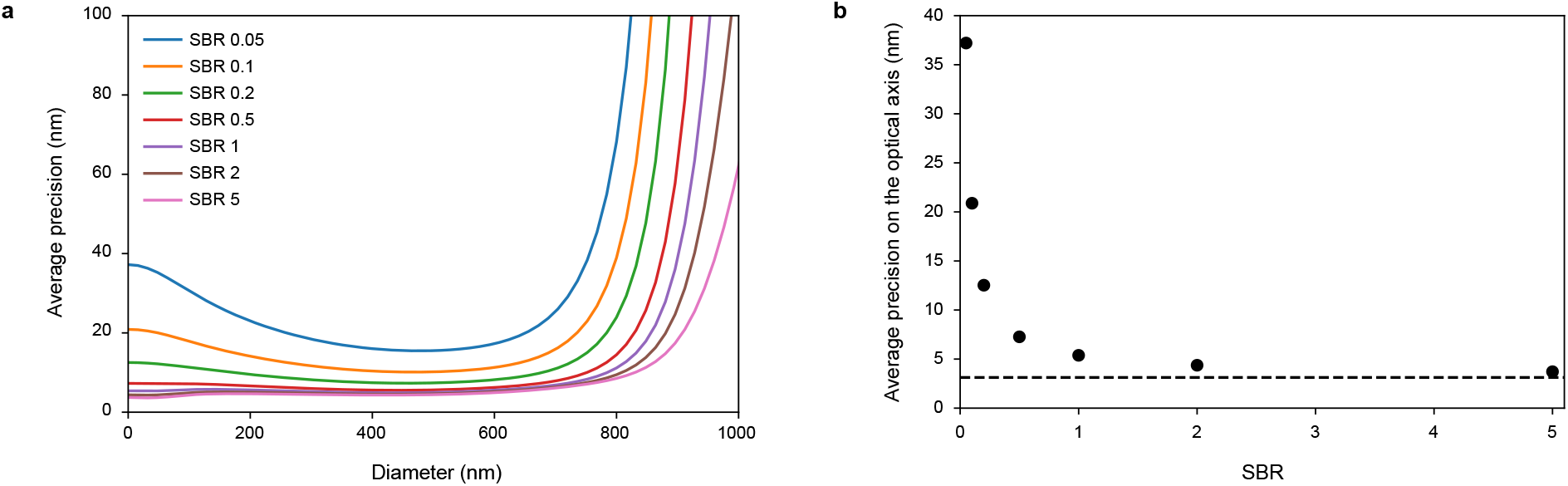
CRB for different SBRs. (a) Average CRB within a circle of a given diameter as a function of the diameter. (b) CRB on the optical axis as a function of the SBR. The dashed line corresponds to the theoretical limit given by Eq. 28. Simulation settings: L = 100 nm, number of emitted photons maximum 1000 in the doughnut maximum, rescaled with the excitation intensity for other positions, corresponding to 135 photons on the optical axis). SBR values are given for the optical axis.

**Figure S6:**
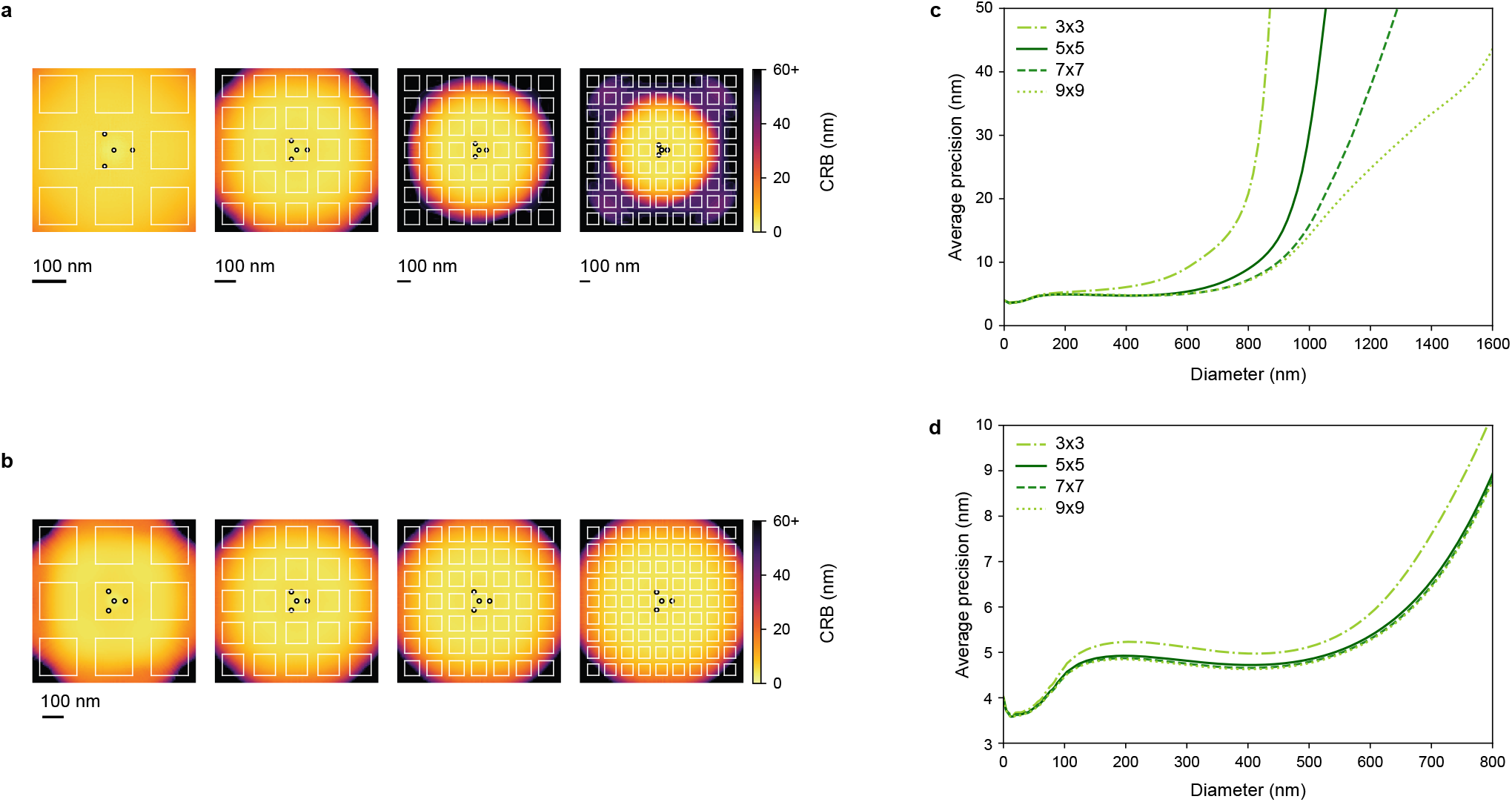
ISM-FLUX CRB for different numbers of detector elements in the array with (a, c) a constant detector element size and (b, d) a constant overall detector size. Simulation settings: *L* = 100 nm, *M* = 500 ×, SBR linearly scaled with the overall excitation intensity, with 100 signal counts and a SBR of 20 for the center.

**Figure S7:**
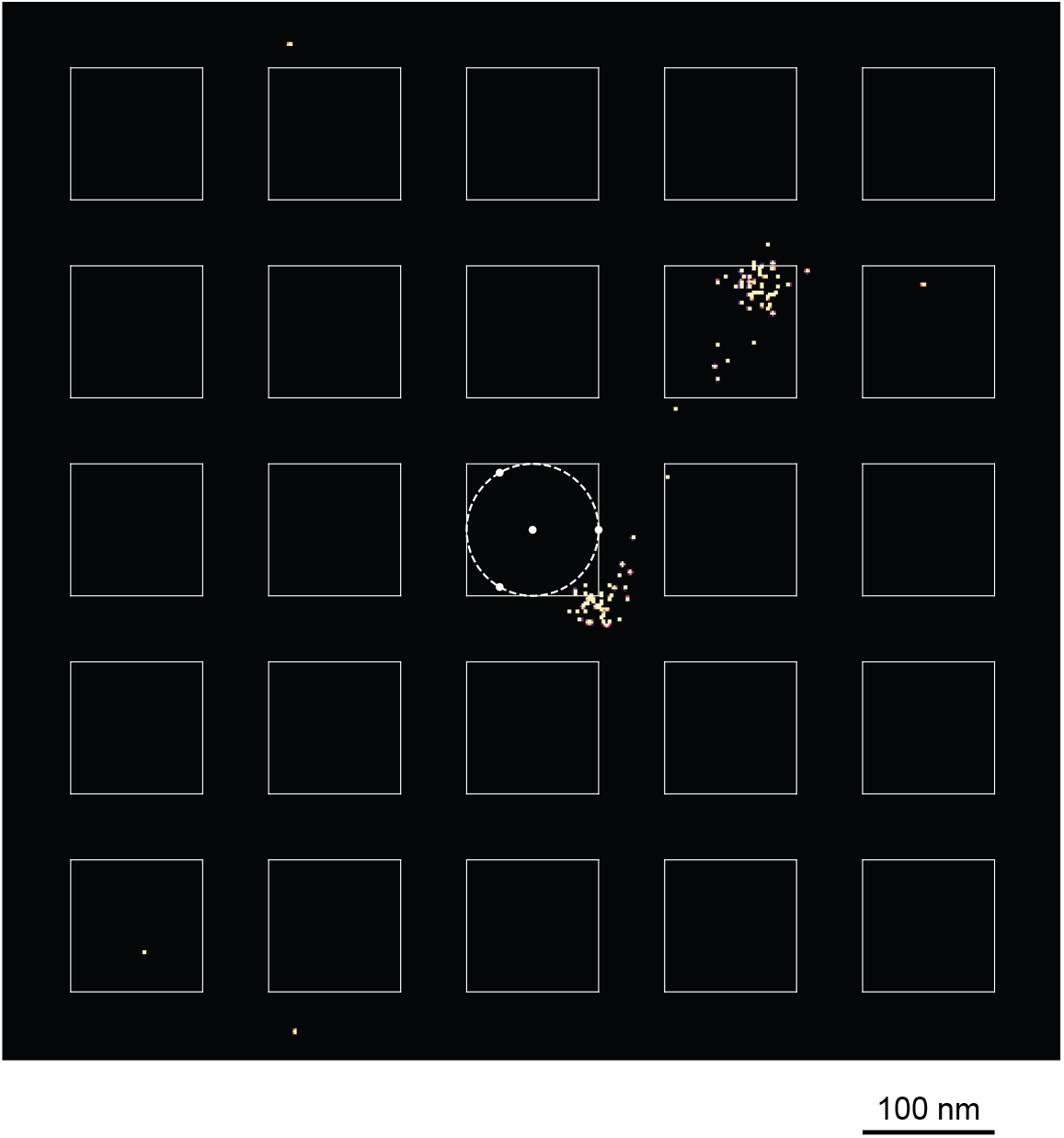
Retrieved localizations for two simultaneously active molecules. L = 100 nm.

**Figure S8:**
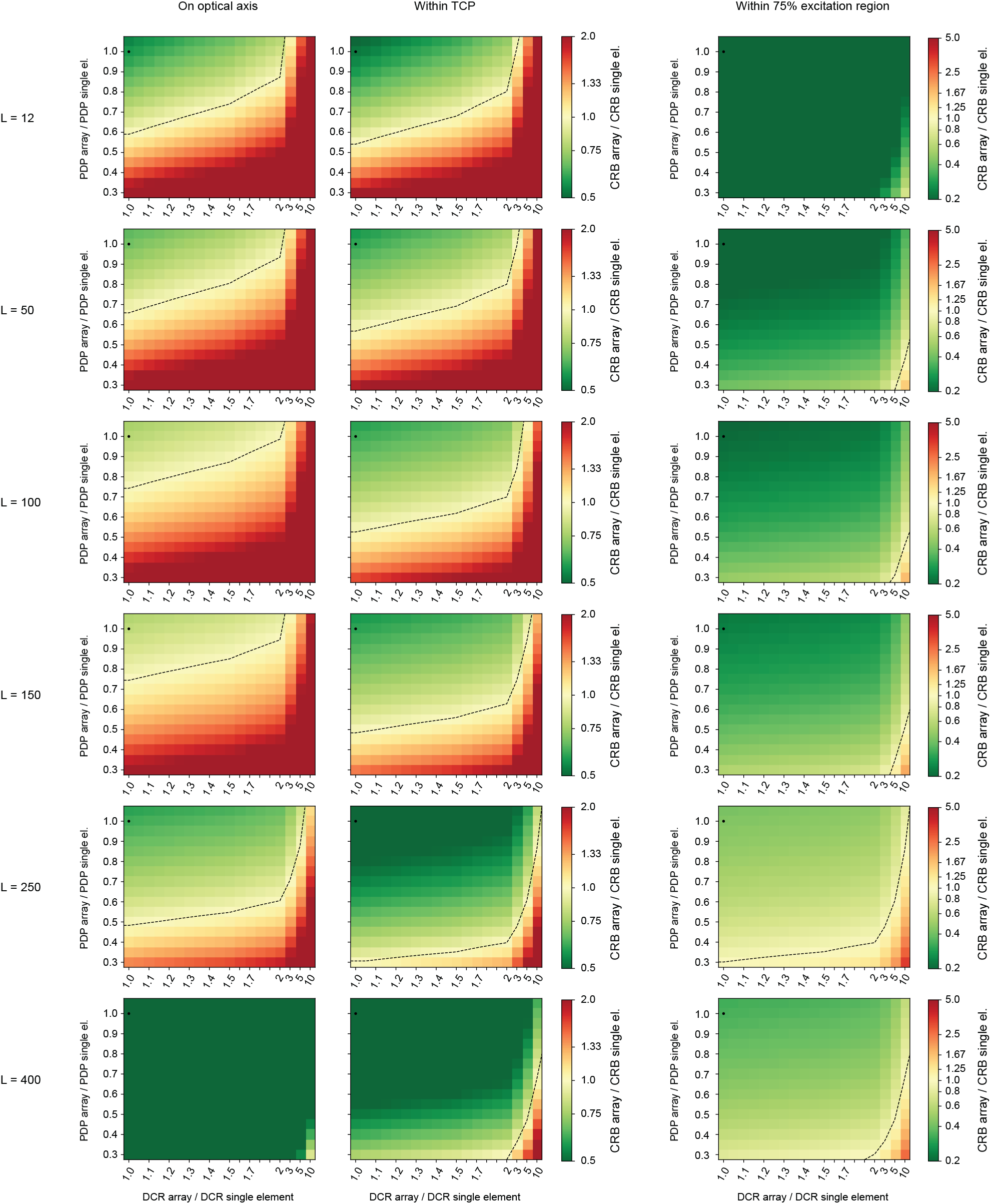
Comparison CRB for localization with an array detector and a single-element detector. Different values of the TCP diameter (*L*, in nm), the photon detection probability (PDP), and the dark count rate (DCR) are shown. Extreme CRB ratios are capped at the color bar limits. The dot in the upper left corner of each panel indicates the data point with coordinates (1, 1), i.e., the situation in which the array detector has the same PDP and DCR as a single-element detector. The dotted lines represent curves of equal performance, separating CRB ratios below from above 1. Simulation settings for the single-element detector: 1000 signal counts for a molecule located in the doughnut maximum, linearly rescaled with the local excitation intensity for the other positions, 125 background counts and a DCR of 125 dark counts. Note that the PDP of the array detector affects both the number of signal counts and the number of background counts. The DCR for the array is defined as the DCR of all pixels combined.

**Figure S9:**
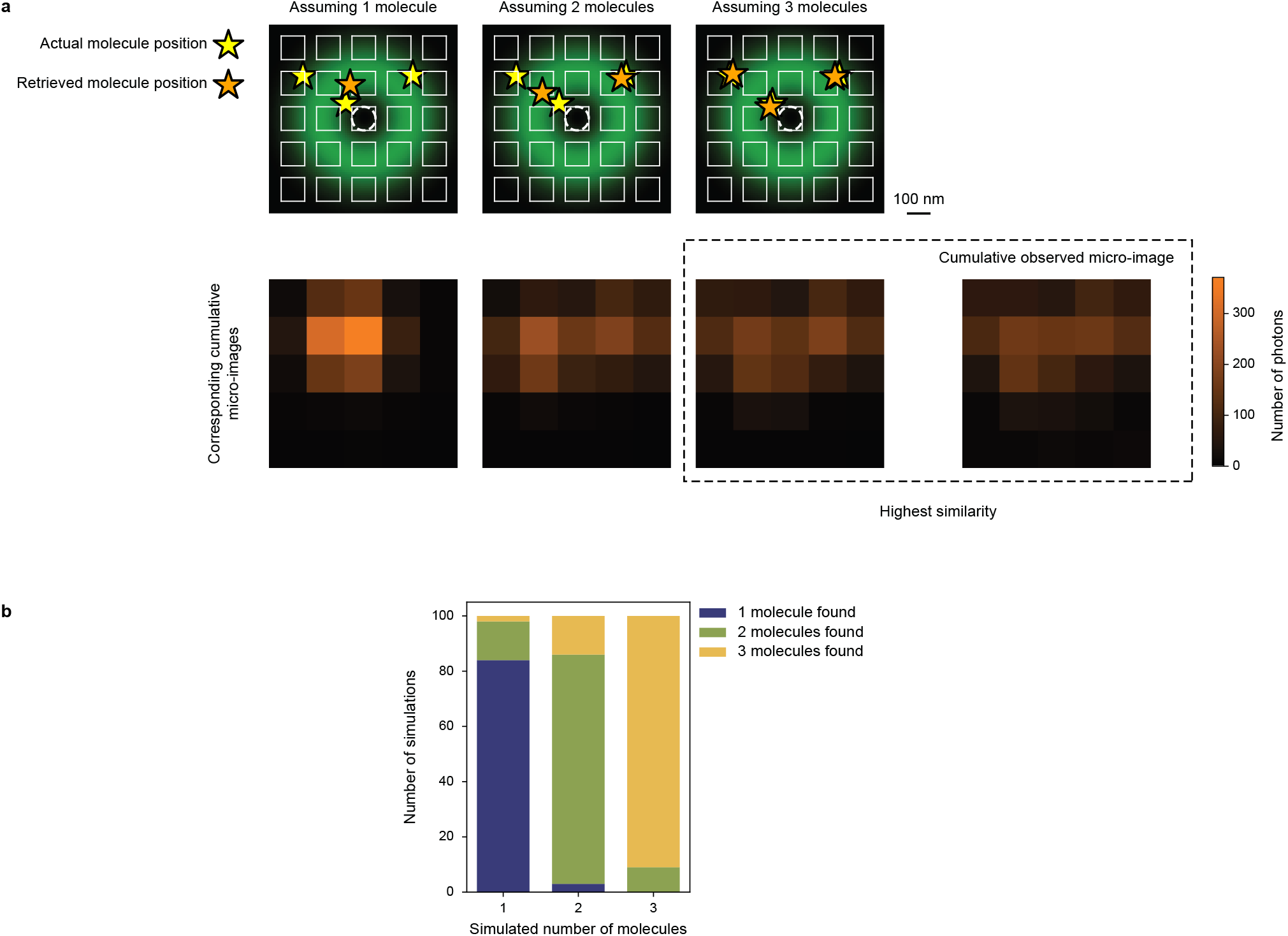
Automatic detection of the number of active fluorophores. (a) Example of the algorithm for three simultaneously active molecules. The algorithm first assumes that only one molecule is active (*N*_*f*_ = 1), tries to localize it (top-left panel), and based on the retrieved position, calculates the expected micro-image for the same number of counts as the experimentally observed micro-image (bottom-left panel). To prevent ending up in a local minimum, the localization step is performed 3 times with each time different starting coordinates, and the result with the lowest negative likelihood is taken. The procedure is repeated for *N*_*f*_ = 2 and *N*_*f*_ = 3 (central and right panels). The resulting micro-images are compared with the experimental micro-image with a similarity metric calculated as the sum of the squared differences between the observed pixel values and the simulated pixel values. The *N*_*f*_ value corresponding to the micro-image with the highest similarity (= smallest difference) is considered to be the correct one. Here, for simplicity, the cumulative micro-images (i.e., the sum over the four excitation patterns) are shown but in reality, the algorithm considers the original 4 × 25 pixel values. An additional filter was applied, removing *subdiffraction* solutions in which two molecules are closer to each other than half a Rayleigh limit. The white circle in the central pixel indicates the TCP. One of the four doughnuts is shown in green. (b) Stacked histograms showing the results for 3 × 100 simulations with molecules randomly placed in the FOV, but within the 75% excitation region and with intermolecular distances of at least half a Rayleigh limit. For every value of *N*_*f*_, the algorithm succeeds to find the actual number of molecules in at least 80% of the cases, with an overall success rate of 86%. Simulation parameters: *L* = 100 nm, 1000 signal counts for each molecule located in the doughnut maximum, linearly rescaled with the local excitation intensity for the other positions, 200 background counts. Counts are for the sum over all detector pixels and all excitation patterns.

**Figure S10:**
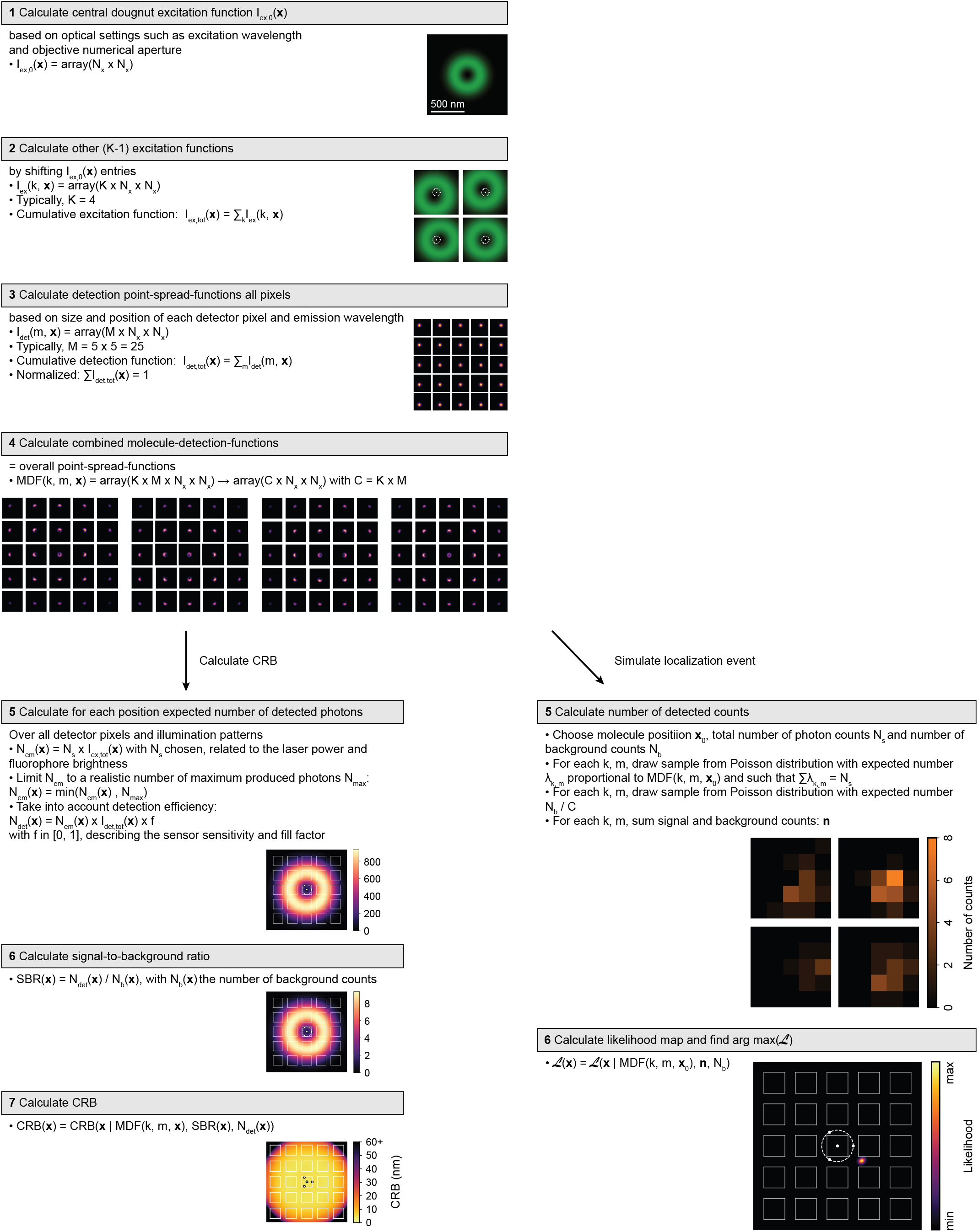
Schematic overview of the simulation protocol.

